# Free fatty acid receptor 4 (FFAR4) regulates cardiac oxylipin balance to promote inflammation resolution in a model of heart failure preserved ejection fraction secondary to metabolic syndrome

**DOI:** 10.1101/2022.04.13.488227

**Authors:** Naixin Zhang, Katherine A. Murphy, Brian Harsch, Michael Zhang, Dylan J. Gyberg, Brandon M. Wagner, Jenna Mendelson, Michael T. Patterson, Devin A. Orchard, Chastity L. Healy, Jesse W. Williams, Gregory C. Shearer, Timothy D. O’Connell

**Author notes:** Correspondence to: Timothy D. O’Connell, PhD, Department of Integrative Biology and Physiology, University of Minnesota School of Medicine, 3-141 CCRB, 2231 6th Street SE, Minneapolis, MN 55414, -and-, Gregory C Shearer, PhD, Department of Nutritional Sciences, 110 Chandlee Laboratory, University Park, PA 16802.

## Abstract

Free fatty acid receptor 4 (Ffar4) is a G-protein coupled receptor for long-chain fatty acids that improves metabolism and attenuates inflammation. Heart failure preserved ejection fraction (HFpEF) is a complex clinical syndrome, but a predominant subset of patients has metabolic syndrome (MetS). Mechanistically, systemic, non-resolving inflammation associated with MetS might promote HFpEF. Interestingly, we recently demonstrated that Ffar4 is cardioprotective in pressure overload. The beneficial effects of Ffar4 on metabolism/inflammation, the high incidence of MetS in HFpEF patients, and the cardioprotective effects of Ffar4 led us to hypothesize that loss of Ffar4 would worsen remodeling in HFpEF secondary to MetS (HFpEF-MetS). To test this, mice with systemic deletion of Ffar4 (Ffar4KO) were fed a high-fat/high-sucrose diet with L-NAME in their water (HFpEF-MetS diet) to induce HFpEF-MetS. In male Ffar4KO mice, the HFpEF-MetS diet induced similar metabolic deficits, but worsened diastolic function and microvascular rarefaction compared to wild-type mice. Conversely, in female Ffar4KO mice, the diet produced greater obesity but no worsening of HFpEF. Loss of Ffar4 in males altered the balance of inflammatory oxylipins in the heart, decreasing the eicosapentaenoic acid derived, pro-resolving oxylipin 18-hydroxyeicosapentaenoic acid (18-HEPE), while increasing the arachadonic acid derived, proinflammatory oxylipin 12-hydroxyeicosatetraenoic acid (12-HETE). This increased 12-HETE/18-HEPE ratio, reflecting a more proinflammatory state, was associated with increased macrophage numbers, which in turn correlated with worsened ventricular remodeling in male Ffar4KO hearts. In summary, our data suggest that Ffar4 controls the pro/anti-inflammatory oxylipin balance in the heart to modulate macrophage function and attenuate HFpEF remodeling.

## INTRODUCTION

Heart failure preserved ejection fraction (HFpEF) is a complex clinical syndrome in which patients present with symptomatic heart failure and a preserved ejection fraction (≥ 50%) (1, 2). Currently in the US, HFpEF prevalence is over 3 million patients, exceeding heart failure reduced ejection fraction (HFrEF), and its incidence is increasing annually (2). Unfortunately, therapies for HFrEF have shown no efficacy in HFpEF (1–3), notwithstanding the recent positive results of empagliflozin to reduce heart failure (HF) hospitalizations in Empagliflozin Outcome Trial in Patients with Chronic Heart Failure with Preserved Ejection Fraction (EMPORER-Preserved) (4). HFpEF etiology is complex and heterogeneous, and patients tend to be older, female, and demonstrate significant phenotypic variation (1, 2). Clinically, attempts have been made to subdivide HFpEF patients based on etiology, indicating that a predominant subset of HFpEF patients has comorbidities associated with metabolic syndrome (MetS) (5–7). Mechanistically, it has been proposed that systemic, non-resolving inflammation associated with MetS and other HFpEF comorbidities may promote HFpEF ventricular remodeling (1, 5, 6), which is characterized by **1)** vascular endothelial cell dysfunction, increased ROS production, and cell death; **2)** inflammation characterized by leukocyte (macrophage) infiltration, leading to activation of fibroblasts and interstitial fibrosis; and **3)** decreased cardiac myocyte nitric oxide (NO) production and dysfunction, all contributing to increased passive stiffness, impaired filling, and exercise intolerance (8–11).

Free fatty acid receptor 4 (Ffar4, GPR120) is a G-protein coupled receptor (GPCR) for endogenous medium- and long-chain saturated, monounsaturated, and polyunsaturated fatty acids (SFA, MUFA, PUFA) (12). This includes, but is not limited to, the cardioprotective ω3-PUFAs, eicosapentaenoic acid (EPA) and docosahexaenoic acid (DHA) (13). Ffar4 is expressed in several tissues with similar expression patterns in mouse and human (14), and high levels of expression in lung, brain, and GI-tract, but lower levels in other tissues including pancreas, small intestine, adipose, taste buds, muscle, heart, liver, and macrophages (14–16).

Ffar4 is a Gq-coupled receptor that activates both Gq- and βarrestin2-mediated pathways and is proposed to improve metabolism and attenuate inflammation (17, 18). However, the role of Ffar4 in regulating metabolism remains somewhat controversial. In general, previous studies in mouse models indicate that in response to a metabolic challenge with high fat diet (HFD), loss of Ffar4 worsens metabolic disease, with evidence of insulin resistance, glucose intolerance, adipose dysfunction, and fatty liver, but with little or no effect on weight gain (19–22). In humans, Ffar4 expression is increased in adipose from obese individuals, and in a European cohort, the Ffar4 R270H inactivating polymorphism is associated with morbid obesity (20). However, other studies have refuted these findings, suggesting loss of Ffar4 does not worsen metabolic disease (23, 24), and in a separate Danish cohort, there was no association between R270H and obesity (25). On the other hand, activation of Ffar4 with synthetic ligands including compound A, TUG-891, or compound 34 generally improves metabolic dysfunction and insulin resistance (26–28).

As noted, Ffar4 also attenuates inflammation in a variety of settings. In macrophages, Ffar4 activation inhibits NFκB signaling and subsequent production of the inflammatory cytokines IL-6 and TNFα, rendering these macrophages less inflammatory and attenuating adipocyte dysfunction in obesity (22, 27). More importantly, Ffar4-mediated activation of cytoplasmic phospholipase A2α (cPLA2α) in macrophages induces the production of oxidatively modified FAs, or oxylipins, that attenuate pro-inflammatory signaling (29–31).

Although we previously detected Ffar4 expression in both cardiac myocytes and fibroblasts (32), little was known about the physiologic function of Ffar4 in the heart until we recently demonstrated that Ffar4 is cardioprotective. Specifically, in male, but not female, Ffar4KO mice following transverse aortic constriction (TAC), we found that Ffar4 is required for an adaptive response to pathologic pressure overload, with worsened systolic and diastolic dysfunction, but surprisingly without excessive fibrosis (33). In adult cardiac myocytes, we observed that Ffar4-cPLA2α signaling specifically and uniquely induced the production of the EPA-derived, cardioprotective, pro-resolving oxylipin 18-hydroxyeicosapentaenoic acid (18-HEPE), which attenuated cardiac myocyte death induced by oxidative stress (33). In humans, we found that *FFAR4* R270H was associated with left ventricular hypertrophy and dilation in a cohort of 7,140 genotyped subjects who had a clinically indicated echocardiogram (33).

The beneficial effects of Ffar4 on metabolism and inflammation (20, 22), the high incidence of MetS in HFpEF patients (7), and the recently described cardioprotective effects of Ffar4 (33) led us to hypothesize that loss of Ffar4 would worsen ventricular remodeling in a mouse model of HFpEF secondary to MetS (HFpEF-MetS). To test this hypothesis, we modified the 2-hit model developed by Schiattarella *et al*. (34) by using a 42% high-fat/30% high-sucrose Western diet, which more closely resembles human dietary patterns, to induce obesity and Type 2 diabetes along with the NO synthase-inhibitor L-NAME in the drinking water to induce hypertension. Here, we report that in male mice with systemic deletion of Ffar4 (Ffar4KO), the HFpEF-MetS diet induced similar metabolic deficits, but worsened diastolic function and microvascular rarefaction relative to wild-type (WT) mice. Conversely, female Ffar4KO mice responded with greater obesity but no worsening of HFpEF. Mechanistically, loss of Ffar4 in males altered the balance of inflammatory oxylipins in the heart, decreasing 18-HEPE levels, while increasing levels of the arachadonic acid (AA) derived, proinflammatory oxylipin 12-hydroxyeicosatetraenoic acid (12-HETE). This increased 12-HETE/18-HEPE ratio, reflecting a more pro-inflammatory state, was associated with increased CD64^+^ macrophages, which in turn correlated with worsened ventricular remodeling in male Ffar4KO hearts. In a broader context, our data suggest that Ffar4 prevents the negative impact of MetS in the heart and suggests Ffar4 might be a novel therapeutic target in cardiometabolic disease.

## METHODS

### Mice

Ffar4KO mice that were generated from cryopreserved sperm from C57Bl/6N-*Ffar4*^tm1(KOMP)Vlcg^ (Design ID: 15078; Project ID: VG15078) purchased from The Mutant Mouse Resource and Research Centers (MMRRC), UC-Davis (Davis, CA, USA). All mice are back-crossed into the C57Bl/6J strain as previously described (33).

At 8 weeks of age, male and female, WT and Ffar4KO mice were randomized to be fed a control diet (DYET #104607, Dyets Inc., Bethlehem, PA) or a diet designed to induce heart failure preserved ejection fraction secondary to metabolic syndrome (HFpEF-MetS) for 20 weeks. Specifically, this HFpEF-MetS diet consisted of the combination of a 42% fat and 30% sucrose chow (DYET #104608, Dyets Inc, Diet composition: Supplemental Tables 1A and 1B) and L-nitroarginine methyl ester (L-NAME, 1 mg/ml, Cat #N5751, Sigma Chemical, St Louis, MO) in the drinking water. Mice were maintained on a 12-hour light/dark cycle at 25°C with *ad libitum* access to food and water. For all experimental analyses, data collection was done with investigator blinded to genotype and treatment.

The HFpEF-MetS diet was designed to induce metabolic syndrome: obesity, hypertension, Type 2 diabetes, increased levels of plasma triglycerides (TG), and decreased levels of plasma high-density lipoprotein (HDL). As such, exclusion criteria included failure to induce obesity (adiposity index (AI) < 0.25) and/or hypertension (systolic blood pressure < 120 mmHg) after 20 weeks on the HFpEF-MetS diet. Of the 86 WT and 93 Ffar4KO mice randomized to the HFpEF-MetS diet, 3 WT male, and 7 WT female were excluded, whereas 5 Ffar4KO male and 2 Ffar4KO female were excluded based on these exclusion criteria.

### Body weight and body composition

Body weights were obtained weekly for all mice for 20 weeks. To measure body composition, EchoMRI scans (EchoMRI, Houston, TX, USA) were performed on all mice after 20 weeks on diet to record fat mass, lean mass, and free water. The adiposity index (AI) was calculated by the equation: AI = fat mass/lean mass.

### Blood pressure

Blood pressure (BP), including mean arterial pressure (MAP), systolic pressure (SP), and diastolic pressure (DP), was measured in all mice after 20 weeks using the CODA non-invasive tail-cuff blood pressure system (Kent Scientific, Torrington, CT, USA). The procedure involved 2 days of acclimation, followed by measurements of blood pressure on 2 subsequent days to determine the average BP. For acclimation, mice were placed in cylindrical restrainers for 15 minutes each day prior to the test. Prior to measurements of BP, the Occlusion (O)-cuff and Volume/Pressure Recording (VPR)-cuff were tested for patency, per manufacturer’s instructions. For measurements of BP, mice were placed into cylindrical restrainers, with the tail left outside to attach the O-cuff and VPR cuff, and placed on a warming plate to acclimate and allow the tail temperature reached 35°C. Once at temperature, the O-cuff was placed at the base of the tail followed by the VPR-cuff. Tail temperature was closely monitored throughout the experiment and maintained between 35-36°C. During each session, 25 blood pressure measurement cycles were recorded, with the first 3 or 4 cycles considered acclimation cycles.

### Intraperitoneal glucose tolerance test

Blood glucose was sequentially measured at 0, 1, 15, 30, 60, and 120 min following an intraperitoneal glucose challenge in all mice after 20 weeks. Briefly, after an overnight fast (14 h), fasting glucose level was measured from tail blood using Quintet AC® Blood Glucose Monitor (McKesson, Irving, TX, USA). Mice were then injected i.p. with D-glucose at 2g/kg fasted body weight (#D16-500, Fisher Scientific, Waltham, MA, USA) and glucose was again measured from tail blood at the indicated time points.

### Triglyceride (TG) and high-density lipoprotein (HDL)

Blood was collected from the retro-orbital plexus into EDTA tubes from all mice after 20 weeks as previously described (33). HDL and TG were measured by colorimetric assays. HDL was measured using a HDL-Cholesterol E kit (Wako Laboratory Chemicals; Richmond, VA) and TG measured using an Infinity reagent colorimetric kit (Thermo Fisher Scientific; Waltham, MA).

### Cardiac function

Cardiac function was measured by echocardiography in all mice after 20 weeks using the Vevo 2100 system (FujiFilm VisualSonics Inc. Toronto, ON, Canada) with a MS400 transducer. For all measurements, mice were anesthetized with isoflurane, gently restrained in the supine position on the prewarmed monitoring pad, and echocardiographic images were captured as mice were recovering from anesthesia to achieve a target heart rate (HR) of 450 – 500 bpm. Isoflurane was maintained at 2-5% and adjusted accordingly in order to maintain a heart rate of 400-500 bpm. Parasternal long axis M-mode images of the left ventricle were captured to measure left ventricular parameters including: left ventricular posterior wall thicknesses (LVPW;s: systolic left ventricular posterior wall; LVPW;d: diastolic left ventricular posterior wall), left ventricular internal diameters (LVID;s: systolic left ventricular internal diameter; LVID;d: diastolic left ventricular internal diameter), left ventricular volumes (ESV: end systolic volume, ((7.0/(2.4 + LVID;s))*LVID;s^3^); EDV: end diastolic volume, ((7.0/(2.4 + LVID;d))*LVID;d^3^)), fractional shortening (FS: 100*((LVID;d – LVID;s)/LVID;d)), ejection fraction (EF: 100*((EDV – ESV)/EDV)), stroke volume (SV: EDV – ESV), and cardiac output (CO: SV*HR).Pulsed-wave Doppler images of the apical four-chamber view and Tissue Doppler images at the level of mitral valve were captured to measure diastolic function, including peak velocity of mitral flow from left ventricular relaxation in early diastole (E wave), peak velocity of mitral flow from left ventricular relaxation in late diastole (A wave), and peak early diastolic mitral annular velocity (e’). Parasternal short axis Pulse-wave Doppler images were captured at the level of aortic valve to measure parameters of pulmonary artery flow including pulmonary acceleration time (PAT) and pulmonary ejection time (PET).

#### Strain Analysis

Global longitudinal strain (GLS) and reverse longitudinal peak strain rate were measured using the speckle-tracking based imaging analysis in the VisualSonics Vevo LAB software version 5.5.0 (Toronto, Canada). Briefly, cine loops of the B-mode in parasternal long axis view were captured. Three consecutive cardiac cycles were chosen to manually draw the endocardial wall border and trace the movement of endocardium over time.

### Tissue histology

Ventricular fibrosis and microvascular density were measured in hearts harvested from mice after 20 weeks. Hearts were injected with 200 μl cold PBS with 60 mM KCl to arrest in diastole, excised, weighed, and subsequently perfused with PBS with 60 mM KCl to clear the heart of blood at 4ml/min rate for 4 mins. Atria were trimmed before hearts were flash-frozen in optimal cutting temperature (OCT) embedding medium. 10-micron cryosections at the midventricular level were cut and mounted at −25°C.

#### Fibrosis

Cryosections were stained in 0.1% solution of Sirius red (direct red 80, Sigma-Aldrich, St Louis, MO) and fast green (Sigma-Aldrich) in 1.2% picric acid (Ricca Chemical Company, Arlington, TX) (20 min, 25°C). To quantify LV fibrotic area, sections were imaged at 10X using a BZ-X800 fluorescent microscope (Keyence, Itasca, IL, USA). Using NIH FIJI software, the color balance of all images was corrected before quantification. Using the color threshold function, picrosirius red positively stained area (fibrotic area) and whole ventricle area (right and left) were defined and fibrotic area/ventricular area was calculated.

#### Microvascular (Capillary) Density

Cryosections were stained with isolectin GS-IB4 (Alexa Fluor™ 594 conjugated; Thermo fisher # I21413) at 20 μg/ml (2 hr, 25°C). and counterstained with DAPI at 1μg/ml (5 min, 25°C). To quantify capillary density, sections were imaged at 20X using a Keyence BZ-X800 fluorescent microscope. Using NIH FIJI software, images were converted to 8-bit images, and following application of the threshold function, images were further converted into bi-color mode. Subsequently, 7-9 regions of interests (ROIs) in a section with a total area between 400000-600000 μm^2^ near the endocardium were randomly selected for analysis. To quantify capillary density, we used an automated quantification algorithm for particle counting (stained capillary endothelium) in which we considered that the diameter of a capillary to be larger than 3 μm, and thus we set the particle size threshold to greater than 7.1 μm^2^. Using this algorithm, average capillary density of the total area was calculated in two representative sections from each heart.

#### Macrophage staining

Cryosections were stained with primary antibodies to detect CD64 (R&D Systems, Minneapolis, MN, USA #AF2074), and subsequently stained with secondary antibodies conjugated to either Alexa fluor 568 (donkey anti-goat, Fisher, Waltham, MA, USA #A11057). To quantify macrophage density, sections were imaged at 20X using a BZ-X800 fluorescent microscope. Using NIH FIJI software, image contrast was adjusted to better visualize the positive staining. Following conversion to 8-bit images, and application of the threshold function, images were further converted into bi-color mode. Subsequently, 7-9 regions of interests (ROIs) with a total area be-tween 600000∼2000000 μm^2^ near the endocardium were randomly selected to calculate the number of CD64^+^ macrophages using an automated quantification algorithm to count particles (positively stained cells). Using this algorithm, macrophage density of the total area was calculated from each heart and was averaged within each treatment group.

### Hydroxyproline content

A hydroxyproline assay kit (Sigma, Burlington, MA, USA #MAK008) was used to quantify total collagen content in the heart. After 20 weeks, hearts were harvested and a portion of the ventricle, ∼20 mg, was frozen and homogenized in ultra-pure water. The ventricular tissue was hydrolyzed in 12M HCl, and after 3 h at 120°C in pressure-tight capped vials, samples were centrifuged at 10000x g for 3 minutes at room temperature. Subsequently, 10 μl (HFpEF samples) or 40 μl (control samples) of supernatant was transferred to the 96-well plate and dried out overnight at 60°C. After drying, the Chloramine T/Oxidation Buffer Mixture provided with the hydroxyproline kit was added to both the sample and standard curve wells and incubated at room temperature. After 5 minutes, diluted DMAB reagent provided with the kit was added to all wells and incubated at 60°C for 90 mins. Finally, absorbance was read at 560nm using a Synergy H1 plate reader (Agilent Technologies, Santa Clara, CA, USA). Hydroxyproline content was calculated using the standard curve and was normalized to tissue weight.

### Cardiac and HDL oxylipin content

Cardiac tissue oxylipins were taken from the apex of mouse hearts. Each tissue sample weight was measured (approximately 50 mg) and recorded and subjected to homogenization followed by oxylipin extraction. Plasma lipoproteins were separated by FPLC followed by measurement of esterified oxylipins in HDL as previously described (REF Murphy et al). HDL and homogenized cardiac tissue samples were spiked with BHT/EDTA (0.2 mg/mL), six deuterated octadecanoid and eicosanoid surrogates (20 µL of 1000 nM concentration with final concentration of 50 nM after reconstitution) and subjected to liquid-liquid extraction to isolate lipid content. Samples were then hydrolyzed in 0.1 M methanolic sodium hydroxide to release ester-linked oxylipins and subjected to solid phase extraction using Chromabond HLB sorbent columns (Machery Nagel, Duren, Germany). Oxylipins were eluted with 0.5 mL of methanol with 0.1% acetic acid and 1 mL of ethyl acetate and dried under nitrogen stream and reconstituted in 200 mL methanol acetonitrile (1:1) with 100 nM of 1-cyclohexyluriedo-3-dodecanoic acid used as internal standard.

Samples were analyzed by liquid chromatography using a Waters Acquity UPLC coupled to Waters Xevo triple quadrupole mass spectrometer equipped with electrospray ionization source (Waters, Milford, MA). 5 mL of the extract was injected, and separation was performed using a CORTECS UPLC C18 2.1 × 100 mm with 1.6 µM particle size column. Flow rate was set at 500 mL/min and consisted of a gradient run using water with 0.1% acetic acid (Solvent A) and acetonitrile isopropanol, 90:10 (Solvent B) for 15 minutes (25% to 95% of Solvent B was set from 0 to 11 min and held at 100% Solvent B from 11 to 13 min. The system was then re-equilibrated and conditioned at 25% Solvent B from 13 to 15 min.). Electrospray ionization operated in negative ion mode with capillary set at 2.7 kV, desolvation temperature set at 600°C, and source temp set to 150°C. Optimal oxylipin MRM transitions were previously identified by direct injection of pure standards onto the mass spectrometer and using cone voltage and collision energy ramps to optimize detection and most prevalent daughter fragments. Calibration curves were generated prior to each run using standards for each oxylipin. Peak detection and integrations were achieved through Target Lynx (Waters, Milford, MA) and each peak inspected for accuracy and corrected when needed.

### RNA isolation and qPCR

RNA was isolated from the apex of the heart using the RNeasy Fibrous Tissue Mini Kit (Qiagen, Germantown MD, USA #74704,) after 20 weeks. RNA concentration was determined by NanoDrop Spectrophotometer (Thermo Fisher, Waltham, MA, USA) and cDNA was synthesized by reverse transcription using the qScript cDNA SuperMix Kit (Quantabio, Beverly, MA, USA #95047-100). Target gene expression was quantified by qRT-PCR using the Bio-Rad (Hercules, CA, USA) iTaq Universal SYBR Green SuperMix (#1725120) and CFX96 Real-Time System.

Primer sequences:

**Ffar4 fwd** CGGCGGGGACCAGGAAAT

**Ffar4 rvs** GTCTTGTTGGGACACTCGGA

**GPR31 fwd** CCACCAGTCTGCCATTCTTTG,

**GPR31 rvs** ACTGTCGTCAGGAAGGCTACT

**ChemR23 fwd** ATGGAGTACGACGCTTACAACG

**ChemR23 rvs** GGTGGCGATGACAATCACCA

### Statistical analysis

Cardiac phenotyping was analyzed using two-way ANOVA with a Tukey’s post-test using Prism 9.0 (GraphPad Software Inc, San Diego, CA). Where specified, principal components analysis (PCA) was used for dimension reduction of oxylipin matrices on log-transformed, standardized concentrations. Mixed models were used to account for within mouse differences in oxylipin concentrations. Statistical significance was set at 0.05; Tukey’s test was used to test for specified post-hoc differences using JMP version 13.2.1.

### Study Approvals

#### Animal

All procedures on animals conformed to the NIH Guide for the Care and Use of Laboratory Animals and were reviewed and approved by the Institutional Animal Care and Use Committee at the University of Minnesota.

#### Additional compliance statements

After 20 weeks, mice were anesthetized with 3% isoflurane, verified by toe-pinch, followed by removal of the heart in accordance with recommendations from the American Veterinary Medical Association. Finally, all the data underlying this article are available in the article and in its online supplementary material.

## RESULTS

### A high-fat/high-sucrose diet with L-NAME (HFpEF-MetS diet) induced similar weight gain, hypertension, and glucose intolerance, but increased triglyceride (TG) and high-density lipoprotein (HDL) levels in male Ffar4KO mice

To test our hypothesis that loss of Ffar4 would worsen HFpEF-MetS, we developed a dietary intervention designed to induce HFpEF secondary to MetS, by adapting the recently described 2-hit model (34). Here, mice were fed a high-fat/high-sucrose diet (42% fat/30% sucrose, less fat and more sucrose than (34)) with L-NAME (1 mg/ml in the drinking water) designed to induce metabolic syndrome (HFpEF-MetS diet). At 8 weeks of age, male and female, WT and Ffar4KO mice were randomized to the control diet (Cont diet: standard low fat diet, no L-NAME) or the high-fat/high-sucrose/L-NAME diet (HFpEF-MetS diet) (Diet composition: Supplemental Tables 1A and 1B, Summary data: Supplemental Table 2A). After 20 weeks, the HFpEF-MetS diet induced similar and significant weight gains in male WT and Ffar4KO mice (Figure 1A). In both WT and Ffar4KO mice, these weight gains were due to significant increases in fat mass (Figure 1B), while lean mass remained unchanged (Figures 1C), resulting in significant increases in the adiposity index (AI, Figure 1D), indicating male WT and Ffar4KO mice developed similar levels of obesity. Furthermore, the HFpEF-MetS diet produced similar and significant increases in mean arterial pressure (MAP) in male WT and Ffar4KO mice (Figure 1E), indicating mild hypertension. The HFpEF-MetS diet also induced a similar degree of glucose intolerance in response to an intraperitoneal glucose challenge in both male WT and Ffar4KO mice (Figure 1F), suggesting progression towards Type 2 diabetes. Interestingly, the HFpEF-MetS diet induced greater increases in both TG and HDL levels in male Ffar4KO mice (Figure 1G, 1H). In summary, 20 weeks of the HFpEF-MetS diet produced a similar weight gain, hypertension, and glucose intolerance, but increased TG and HDL levels in male Ffar4KO mice relative to WT mice.

**Figure 1.**
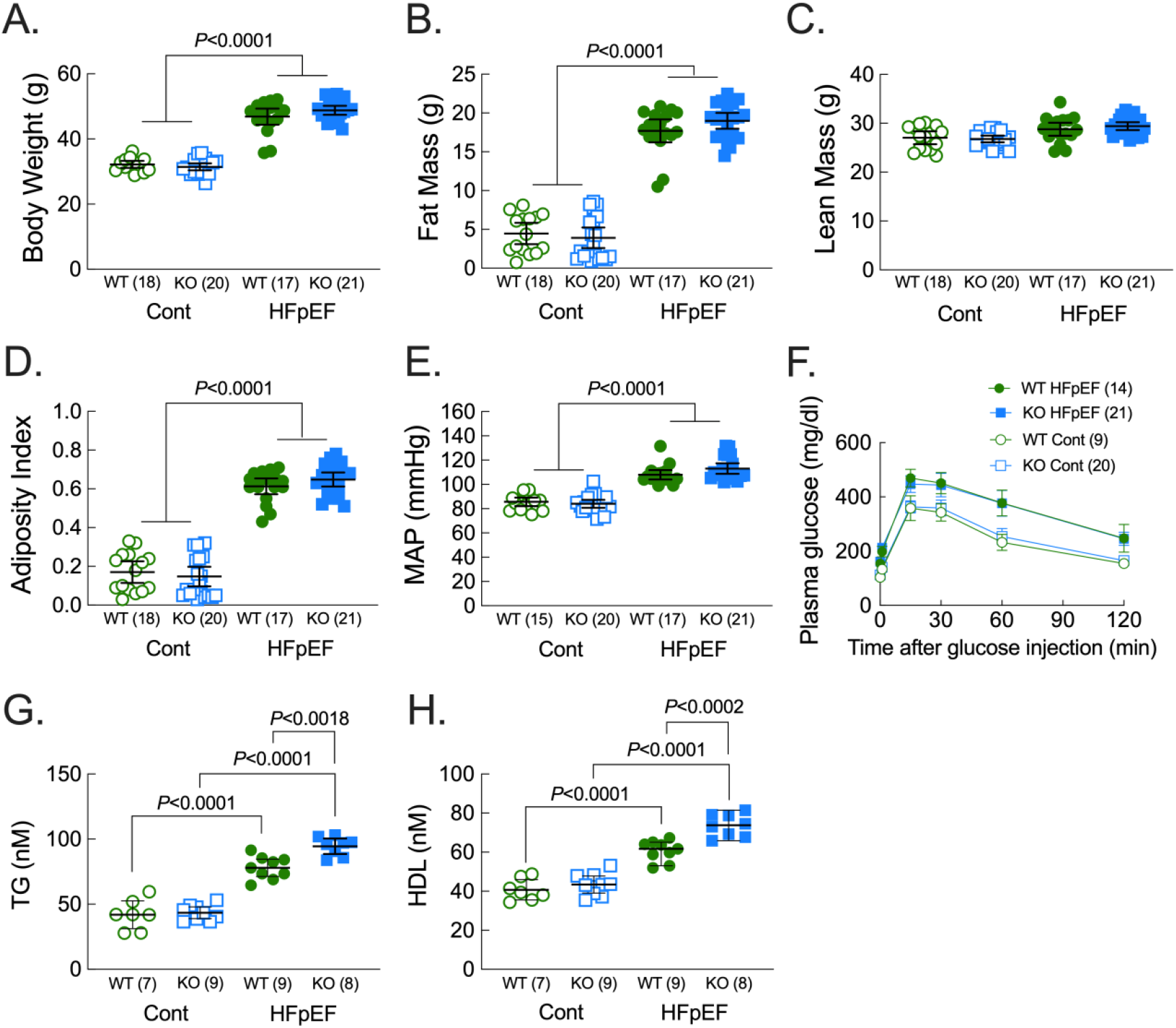
A high-fat/high-sucrose diet with L-NAME (HFpEF-MetS diet) induced similar weight gain, hypertension, and glucose intolerance, but increased triglyceride (TG) and high-density lipoprotein (HDL) levels in male Ffar4KO mice. Male wild-type (WT, green) and Ffar4KO (KO, blue) mice were fed a control diet (Cont., open symbols) or the combination of a high-fat (42%)/high-sucrose (30%) diet and L-NAME (1 mg/ml) in the drinking water (HFpEF, closed symbols) for 20 wks. After 20 weeks; **A**. body weight was recorded, and body composition, including: **B**. Fat Mass and **C**. Lean Mass were determined by EchoMRI; and **D**. Adiposity Index (fat mass/lean mass) was calculated. **E**. Mean Arterial Pressure (MAP) measured by tail-cuff. **F**. [Intraperitoneal glucose tolerance test](https://www.mousephenotype.org/impress/ProcedureInfo?action=list&procID=399) (IPGTT). **G**. Triglyceride levels and **H**. High density lipoprotein levels were measured from plasma. Data (**A-H**) are presented as Mean ± 95% CI and were analyzed by two-way ANOVA with Tukey’s multiple comparison test.

### Metabolic syndrome worsened diastolic dysfunction and microvascular rarefaction in male Ffar4KO mice

Previous studies in mice (34, 35) and humans (7) have demonstrated a link between MetS and HFpEF. After 20 weeks, we measured cardiac function by echocardiography in both male WT and Ffar4KO mice to determine if MetS induced HFpEF in WT mice and if loss of Ffar4 worsened cardiac function secondary to MetS (Figure 2, Supplemental Table 3A). In agreement with previous studies (34, 35), the HFpEF-MetS diet induced significant diastolic dysfunction, evidenced by increased E/e’ ratio and E/A ratios (Figure 2A-B, green symbols), but ejection fraction was preserved (EF, Figure 2C, green symbols) in male WT mice. More importantly, in male Ffar4KO mice, the HFpEF-MetS diet significantly worsened diastolic dysfunction relative to male WT mice (E/e’, E/A, Figure 2A-B, closed green symbols vs closed blue symbols), but again EF was preserved (Figure 2C). Furthermore, the HFpEF-MetS diet decreased global longitudinal strain (GLS, Figure 2D), and reduced reverse longitudinal peak strain rate (Figure 2E) in male Ffar4KO mice, the latter indicating worsened diastolic function (36–38). HFpEF is also associated with left atrial dysfunction and pulmonary hypertension (39, 40), and we measured pulmonary acceleration time (PAT) in both male WT and Ffar4KO mice as an indicator of pulmonary arterial pressure (41, 42). Interestingly, the HFpEF-MetS diet induced a significant drop in PAT only in male Ffar4KO mice relative to Ffar4KO mice on the control diet, with no effect in male WT mice (Figure 2F), suggestive of pulmonary hypertension in the male Ffar4KO mice.

**Figure 2.**
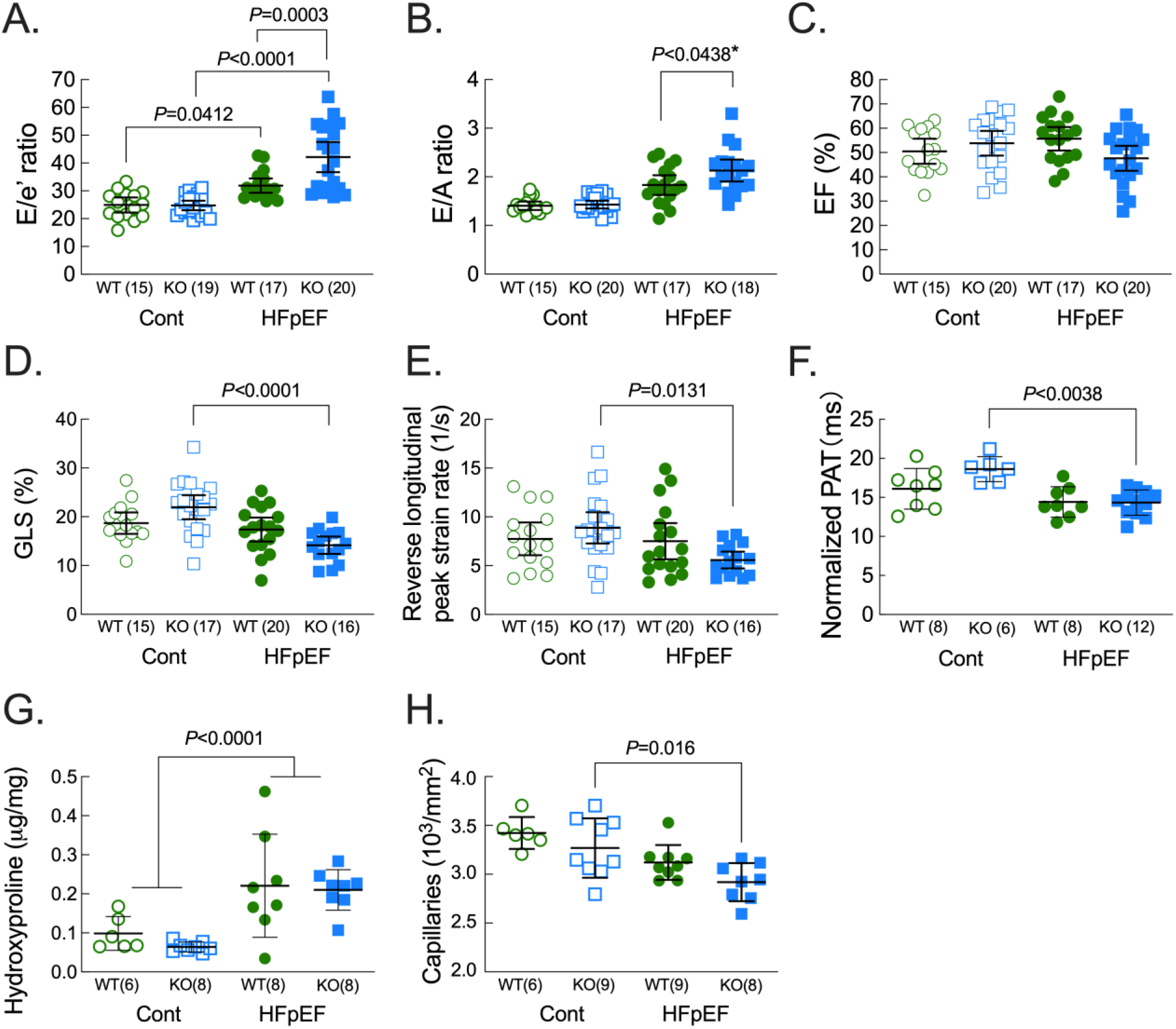
Metabolic syndrome worsened diastolic dysfunction and microvascular rarefaction in male Ffar4KO mice. Cardiac function was measured by echocardiography in male wild-type (WT, green) and Ffar4KO (KO, blue) mice after 20 weeks on the control diet (Cont., open symbols) or HFpEF diet (HFpEF, closed symbols). **A**. E/e’ ratio, **B**. E/A ratio, **C**. ejection fraction (EF,%), **D**. global longitudinal strain (GLS, %) **E**. reverse longitudinal peak strain rate (1/s) **F**. pulmonary acceleration time (PAT, ms) normalized by heart rate from male WT and Ffar4KO mice. **G**. Ventricular fibrosis quantified hydroxyproline content (μg/mg ventricular wet weight). **H**. LV myocardial capillary density quantified by Isolectin-B4. Data (**A-H**) are presented as Mean ± 95% CI and were analyzed by two-way ANOVA with Tukey’s multiple comparison test. * Primary interaction: *P*=0.0779, Welch’s t-test for WT HFpEF vs Ffar4KO HFpEF, employed due to unequal variance between Control and HFpEF diet groups: *P*=0.0438.

Two prominent changes to cardiac structure associated with HFpEF are an increase in interstitial fibrosis and microvascular rarefaction (11). After 20 weeks, total cardiac collagen content, measured by hydroxyproline content, was significantly increased to a similar degree in both male WT and Ffar4KO mice (Figure 2G). Interestingly, there was no detectable change in interstitial fibrosis measured by picrosirius red staining, which stains primarily fibrillar collagens I and III, in either male WT or Ffar4KO mice (Supplemental Figure 1B), similar to the very minor changes previously observed in a similar model (34). However, a failure to observe interstitial fibrosis with picrosirius red staining was noted in a mouse model of chronic kidney disease-induced HFpEF, although a further proteomic analysis revealed profound changes in the extracellular matrix, including many non-fibrillar collagen (43), suggesting the failure to observe an increase in interstitial fibrosis could be a technical issue. Furthermore, the HFpEF-MetS diet reduced capillary density in male Ffar4KO mice relative to Ffar4KO mice on the control diet (Figure 2H). In summary, the loss of Ffar4 in males despite having little to no effect on MetS, significantly worsened diastolic function and microvascular rarefaction, suggesting that Ffar4 is required for an adaptive response to pathological cardiovascular stress induced secondary to MetS in males.

### The HFpEF-MetS diet induced greater weight gain but no worsening of diastolic function in female Ffar4 KO mice

In female WT and Ffar4KO mice, the HFpEF-MetS diet induced significantly greater weight gain in the Ffar4KO mice (Figure 3A, Summary data: Supplemental Table 2B). This additional weight gain in female Ffar4KO mice was due to a significant increase in fat mass, while lean mass remained unchanged, resulting in a significant increase in the AI (Figure 3B), indicating a greater degree of obesity in the female Ffar4KO mice. However, the HFpEF-MetS diet produced similar degrees of hypertension and similar increases in TG and HDL levels in female WT and Ffar4KO (Figure 3C, 3E, 3F), but interestingly failed to induce glucose intolerance in either WT or Ffar4KO mice (Figure 3D). In female WT mice, the HFpEF-MetS diet induced significant diastolic dysfunction evidenced by increased E/e’ and E/A ratios (Figures 3G-H, orange symbols, Summary data: Supplemental Table 3B) with preserved EF (Figure 3I, orange symbols). However, despite the increased obesity observed in the female Ffar4KO mice, the HFpEF-MetS diet did not result in any further worsening of diastolic function relative to female WT mice, and EF was preserved (Figures 3G-I, closed orange versus closed purple symbols). In summary, the loss of Ffar4 in females resulted in greater obesity in response to the HFpEF-MetS diet, but this had no worsening effect on cardiac function as was observed in male Ffar4KO mice. This sex-based difference was similar to our prior report (33), and as such, we did not investigate the cardiac phenotype in females further.

**Figure 3.**
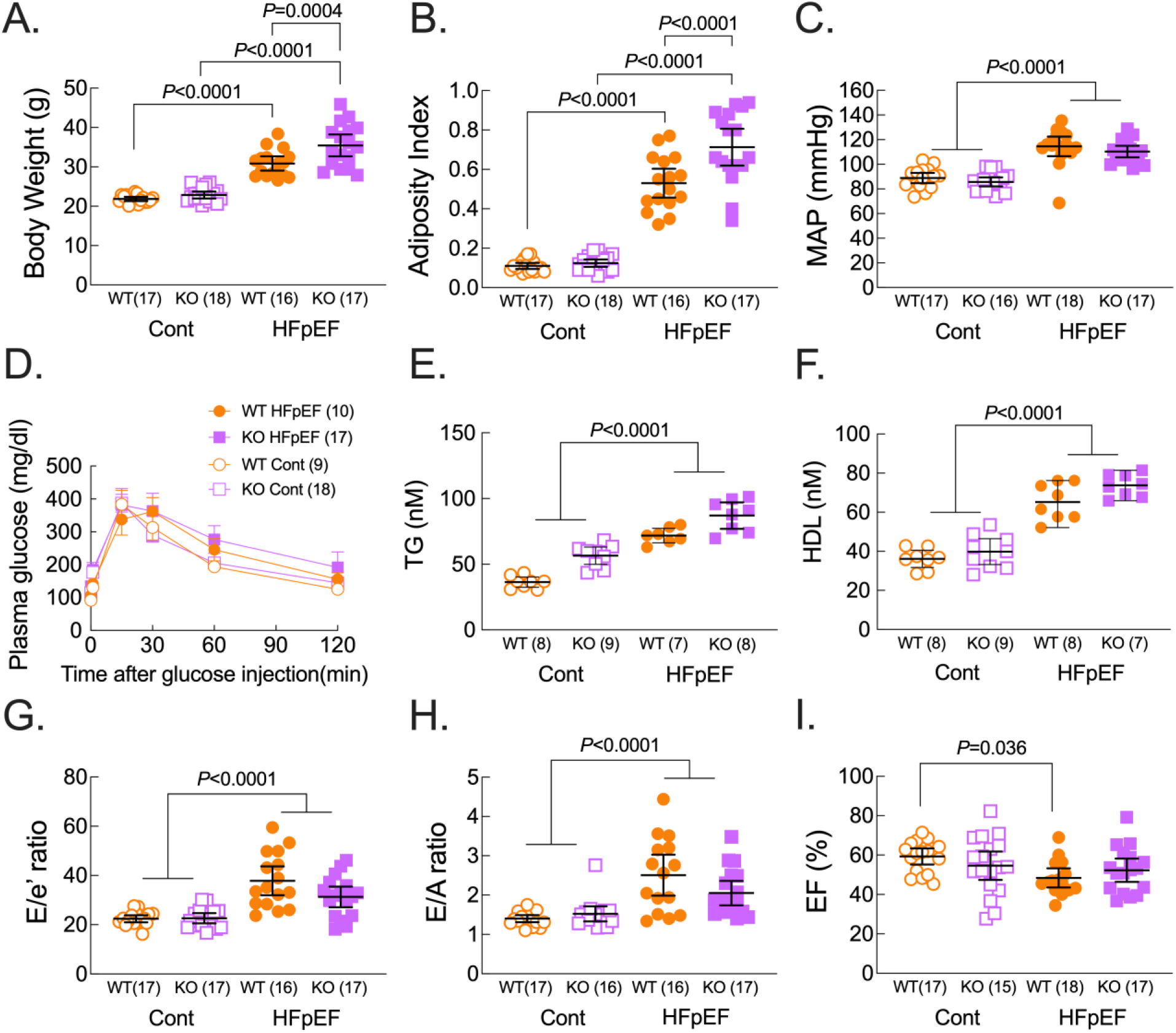
The HFpEF-MetS diet induced greater weight gain but no worsening of diastolic function in female Ffar4 KO mice. Female wild-type (WT, orange) and Ffar4KO (KO, purple) mice were fed a control diet (Cont., open symbols) or the combination of a high-fat (42%)/high-sucrose (30%) diet and L-NAME (1 mg/ml) in the drinking water (HFpEF, closed symbols) for 20 wks. After 20 weeks; **A**. body weight was recorded and **B**. Adiposity Index (fat mass/lean mass, determined by EchoMRI) was calculated. **C**. Mean Arterial Pressure (MAP) measured by tail-cuff. **D**. [Intraperitoneal glucose tolerance test](https://www.mousephenotype.org/impress/ProcedureInfo?action=list&procID=399) (IPGTT). **E**. Triglyceride levels and **F**. High density lipoprotein levels were measure from plasma. Cardiac function was measured by echocardiography. **G**. E/e’ ratio, **H**. E/A ratio, **I**. ejection fraction (EF,%). Data (**A-I**) are presented as Mean ± 95% CI and were analyzed by two-way ANOVA with Tukey’s multiple comparison test.

### Metabolic syndrome decreased 18-HEPE, a pro-resolving EPA-derived oxylipin, and increased 12-HETE, a pro-inflammatory AA-derived oxylipin, in HDL and hearts of male Ffar4KO mice

We previously demonstrated that in cardiac myocytes, Ffar4-cPLA2α signaling specifically induced the production of the EPA-derived, pro-resolving oxylipin 18-HEPE, which protected cardiac myocytes from oxidative stress (33). Interestingly, a recent report indicated that the arachadonic acid (AA)-derived, pro-inflammatory oxylipin 12-hydroxyeicosatetraenoic acid (12-HETE) exacerbates endothelial dysfunction, and that antagonizing 12-HETE improved outcomes in mouse model of HFpEF in Type 2 diabetic db/db mice (44). Here, we investigated the relationship between 18-HEPE and 12-HETE in HFpEF secondary to MetS and how loss of Ffar4 might impact 18-HEPE and 12-HETE levels, which we measured in circulating HDL (representing oxylipins exported from cells in the heart and other tissues) and in both the esterified (membrane) and non-esterified (NEOx, free) fractions in the heart (Figure 4).

**Figure 4.**
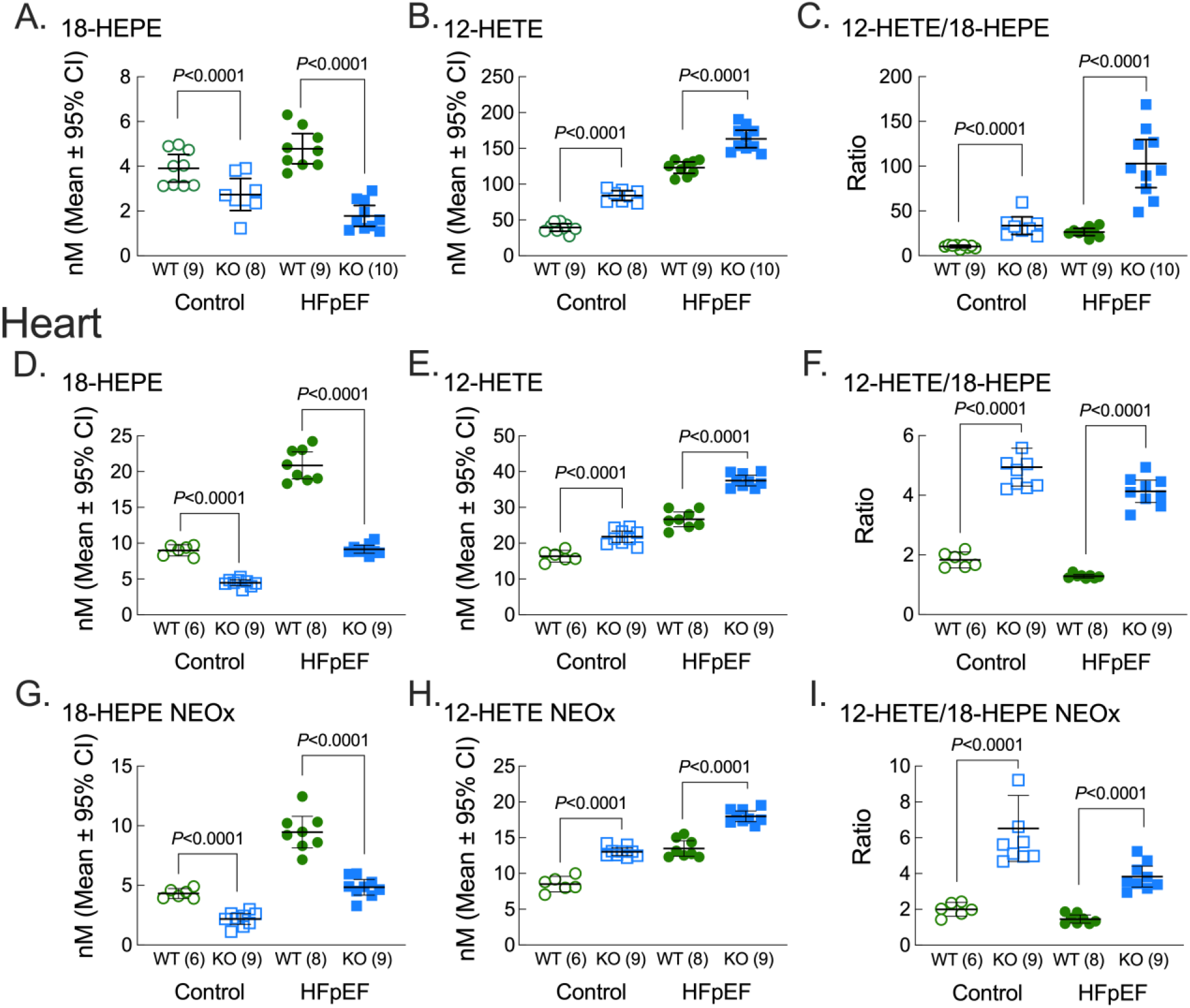
Metabolic syndrome decreased 18-HEPE, a pro-resolving EPA-derived oxylipin, and increased 12-HETE, a pro-inflammatory AA-derived oxylipin, in HDL and hearts of male Ffar4KO mice. **A-C**. After 20 weeks on diet, plasma was collected and HDL oxylipin content was detected by liquid chromatography/mass spectrometry. Levels of the **A**. 18-HEPE, an EPA-derived, pro-resolving oxylipin; **B**. 12-HETE, an AA-derived, pro-inflammatory oxylipin; and **C**. 12-HETE/18-HEPE ratio are shown. **D-I**. In a similar fashion, after 20 weeks, hearts were collected and oxylipin content, esterified and non-esterified (NEOx), were detected by liquid chromatography/mass spectrometry. **D, G**. 18-HEPE, esterified, non-esterified; **E, H**. 12-HETE, esterified, non-esterified; and **F, I**. 12-HETE/18-HEPE ratio, esterified, non-esterified. Data are presented as Mean ± 95% CI and were analyzed by two-way ANOVA with Tukey’s multiple comparison test.

#### WT mice

In male WT mice, the HFpEF-MetS diet significantly increased both 18-HEPE and 12-HETE levels in HDL (Figure 4A-B, green symbols) and both were also significantly increased in the esterified (Figures 4D-E) and non-esterified fractions in the heart (Figures 4G-H). As a result, in male WT mice fed the HFpEF-MetS diet, the 12-HETE to 18-HEPE ratio remained relatively constant (Figure 4C, F, I, green symbols), with only a relatively small increase in HDL and a slight decrease in both the esterified (Figure 4F) and non-esterified (Figure 4I) fractions in the heart.

#### Ffar4KO mice

In male Ffar4KO mice, 18-HEPE levels were significantly lower in HDL and the esterified and non-esterified fraction in the heart relative to WT mice on the control diet (Figures 4A, D, G, respectively, compare open symbols), consistent with our prior report that Ffar4 is largely responsible for 18-HEPE synthesis (33). Unlike in male WT mice, the HFpEF-MetS diet decreased 18-HEPE levels in HDL, and induced smaller increases in 18-HEPE in the esterified and non-esterified fractions in the heart in male Ffar4KO mice (Figures 4A, D, G, compare blue symbols). Conversely, 12-HETE levels were elevated in HDL and the esterified and non-esterified fractions in the heart from Ffar4KO mice relative to WT mice on the control diet (Figures 4B, E, H, respectively, compare open symbols). Furthermore, the HFpEF-MetS diet increased 12-HETE levels in all fractions in the Ffar4KO mice (Figures 4B, E, H, respectively, compare blue symbols), and the absolute levels of 12-HETE in all fractions in Ffar4KO mice were greater than in WT mice (Figures 4B, E, H, respectively, compare closed symbols). As a result, in male Ffar4KO mice, the 12-HETE to 18-HEPE ratio was slightly increased in HDL, with a larger increase in the esterified and non-esterified fraction in the heart in mice on the control diet (Figures 4C, F, I, respectively, compare open symbols). Furthermore, in male Ffar4KO mice fed the HFpEF-MetS diet, the 12-HETE to 18-HEPE ratio was dramatically increased in the HDL and showed a similar increase relative to WT mice on the control diet in the esterified and non-esterified fractions (Figures C, F, I, respectively, compare the closed symbols).

In summary, these results reveal some generalized trends in Ffar4-dependent oxylipin synthesis. First, the loss of Ffar4 is correlated with decreased synthesis of 18-HEPE, but increased synthesis of 12-HETE. Second, MetS is associated with increased 18-HEPE and 12-HETE synthesis, such that the overall balance between the pro-inflammatory 12-HETE and proresolving 18-HEPE is largely maintained. Finally, and most importantly, the loss of Ffar4 dramatically impairs the ability to increase 18-HEPE synthesis while further increasing 12-HETE synthesis. The net result being that in the Ffar4KO, the 12-HETE to 18-HEPE ratio is significantly increased, suggesting a more pro-inflammatory state in the male Ffar4KO heart that correlated with the worsened cardiovascular outcomes in the Ffar4KO in response to the HFpEF-MetS diet.

### Metabolic syndrome increased the expression of Ffar4 in male WT hearts, as well as ChemR23, a receptor for E-series Resolvins, in male Ffar4KO hearts

18-HEPE and 12-HETE are agonists that can lead to downstream activation of GPCR signaling, 18-HEPE through conversion to E-Series resolvins (RvE1) and activation of ChemR23 (45), and 12-HETE through activation of GPR31 (46) and BLT1 receptors (47). Therefore, to define the relationship between the increased levels of 18-HEPE, decreased levels of 12-HETE in Ffar4KO mice, and expression of their cognate receptors in the heart, we measured whole heart Ffar4, ChemR23, and GPR31 mRNA expression. In WT mice, the HFpEF-MetS diet significantly increased Ffar4 expression (Figure 5A), a somewhat surprising finding given that we previously found that Ffar4 expression is decreased in human HF (33). In Ffar4KO mice, the HFpEF-MetS diet significantly increased ChemR23 expression relative to WT (Figure 5B), while GPR31 expression was elevated in the Ffar4KO heart regardless of diet (Figure 5C). The increased levels of ChemR23 and GPR31 in the male Ffar4KO heart, along with the changes in their respective ligands support the idea that the HFpEF-MetS diet might have induced a more pro-inflammatory state in the male Ffar4KO heart.

**Figure 5.**
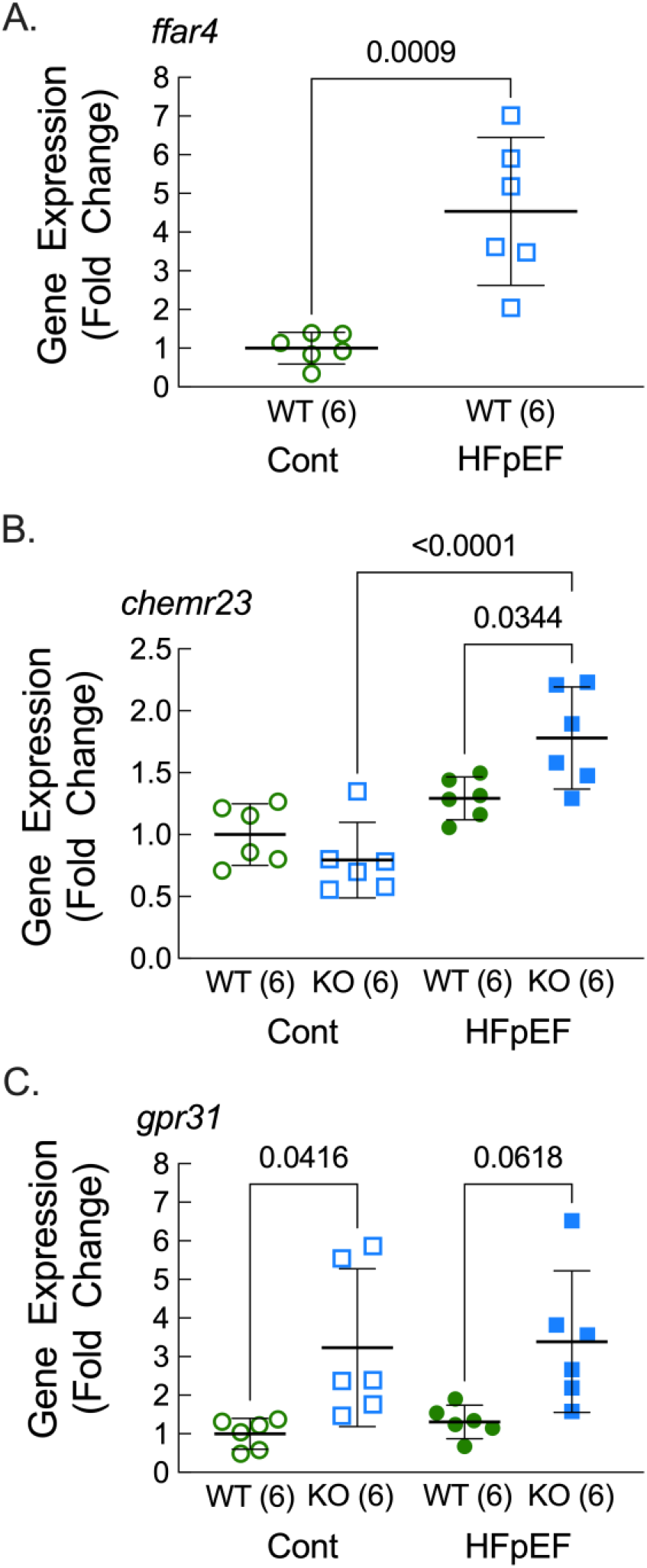
Metabolic syndrome increased the expression of Ffar4 in male WT hearts, as well as ChemR23, a receptor for E-series Resolvins, in male Ffar4KO hearts. **A-C**. After 20 weeks on diet, RNA was isolated and **A**. Ffar4, **B**. ChemR23 and **C**. GPR31 mRNA levels were quantified by qRT-PCR. Data are presented as Mean ± 95% CI and were analyzed by two-way ANOVA with Tukey’s multiple comparison test.

### Metabolic syndrome increased CD64^+^ macrophages in male Ffar4KO hearts, which correlated with worsened ventricular remodeling

Recently, it has become clear that cardiac macrophages represent a diverse cell population that plays a critical role in cardiac homeostasis and the response to cardiac injury (Reviews: (48, 49)). Furthermore, systemic inflammation and recruitment of macrophages to the heart might promote HFpEF, and a few studies have suggested that macrophage numbers are increased in models of hypertension/chronic kidney disease or aging that resemble HFpEF (50–52), as well as human HFpEF patients (52). With the increase in the 12-HETE/18-HEPE ratio in the male Ffar4KO mice suggesting a more inflammatory state in the heart, we hypothesized that macrophage numbers would be increased in Ffar4KO hearts. To test this hypothesis, we quantified the number of CD64^+^ macrophages in male WT and Ffar4KO hearts. In male WT hearts, the HFpEF-MetS diet increased the number of CD64^+^ macrophages, indicating an increase in total macrophage number, while the HFpEF-MetS diet induced a further significant increase in CD64^+^ macrophages in the male Ffar4KO hearts (Figure 6A, B). The increase CD64^+^ macrophages in WT hearts in response to MetS supports the hypothesis that inflammation drives HFpEF remodeling. More importantly, the further increase in this CD64^+^ macrophage population in Ffar4KO hearts in response to MetS is consistent with the increased 12-HETE/18-HEPE ratio in these mice. Interestingly, the number of CD64^+^ macrophages was positively correlated with diastolic dysfunction (E/e’, Figure 6C, E/A, Figure 6D) and inversely correlated with microvascular rarefaction (Figure 6E), again supporting the hypothesis that inflammation drives HFpEF remodeling.

**Figure 6.**
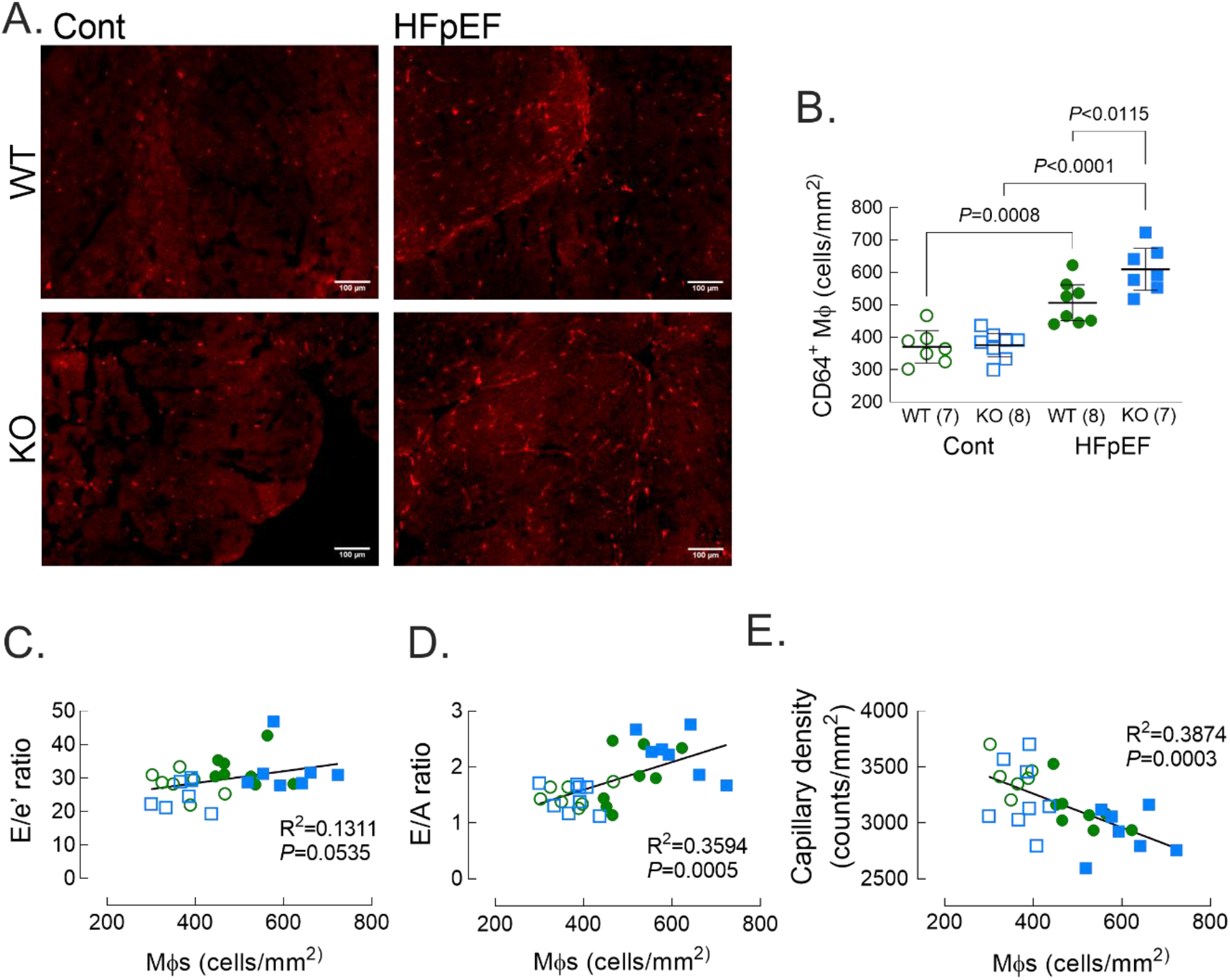
Metabolic syndrome increased CD64^+^ macrophages in male Ffar4KO hearts, which correlated with worsened ventricular remodeling. **A-B**. After 20 weeks on diet, ventricular sections were stained CD64 to detect the total macrophage population (**A**) and total CD64^+^ cells were counted (**B**). Data (**B**) are presented as Mean ± 95% CI and were analyzed by two-way ANOVA with Tukey’s multiple comparison test. **C-E**. Total CD64^+^ macrophages were correlated with markers of HFpEF remodeling: **C**. E/e’ ratio; **D**. E/A ratio; and **E**. Capillary density.

## DISCUSSION

Of the roughly 6 million total cases of HF in the US, the prevalence of HFpEF now exceeds 50% (53), and despite the recent success of empagliflozin in EMPORER-Preserved (4), therapeutic options for HFpEF remain limited. Here, we have identified a novel cardioprotective role for Ffar4 in the context of HFpEF secondary to MetS, identifying a potentially new therapeutic target for the management of cardiometabolic disease. In response to a dietary challenge designed to induce MetS, modified from the original 2-hit HFpEF model proposed by Schiattarella *et al*. (34), we found that systemic deletion of Ffar4 surprisingly did not affect MetS, but did significantly worsen HFpEF remodeling in male Ffar4KO mice. Conversely, the HFpEF-MetS diet induced greater obesity, but no worsening of HFpEF remodeling in female Ffar4KO mice. Interestingly, we found that levels of the EPA-derived, cardioprotective oxylipin 18-HEPE were significantly decreased in HDL and hearts from male Ffar4KO mice. On the contrary, levels of 12-HETE, a pro-inflammatory, AA-derived oxylipin, were significantly increased in HDL and hearts from male Ffar4KO mice. This resulted in an increase in the pro-inflammatory 12-HETE to pro-resolving 18-HEPE ratio in male Ffar4KO mice on the HFpEF-MetS diet, suggesting a more pro-inflammatory state that correlated with the worsened HFpEF remodeling in these mice. 18-HEPE is the sole precursor for E-series resolvins (RvE), which are ligands for the GPCR ChemR23 (45), whereas 12-HETE is a ligand for GPR31 (46) and leukotriene B4 receptor BLT1 (47). In male Ffar4KO mice, ChemR23 expression was lower at baseline, but showed a much larger increase in response to the HFpEF-MetS diet, while GPR31 was elevated regardless of diet in the male Ffar4KO, suggesting a potential link between decreased 18-HEPE levels, increased 12-HETE levels, and macrophage function in male Ffar4KO mice on the HFpEF-MetS diet. As mentioned, HFpEF is proposed to be a disease of systemic inflammation (1), and in fact, we provide the first evidence in this 2-hit HFpEF-MetS model that CD64^+^ macrophage numbers are increased in WT hearts. More importantly, we observed a further 2-fold increase in CD64^+^ macrophages in male Ffar4KO hearts, correlating with the heightened pro-inflammatory state in these mice. Briefly, the data suggest that Ffar4 controls the pro/anti-inflammatory oxylipin balance in the heart to modulate macrophage function and attenuate HFpEF remodeling.

Inflammation secondary to comorbidities associated with HFpEF, including MetS and chronic kidney disease, are proposed to drive HFpEF remodeling (1, 5, 6). In support of this hypothesis, inflammation associated with hypertension, obesity, and diabetes predicted incident HFpEF, but not HFrEF in the Health, Aging, and Body Composition study (Health ABC) (54). Plasma markers of inflammation including soluble IL-1 receptor like 1 (IL1RL1), C-reactive protein (CRP), growth and differentiation factor 15 (GDF15) are high in HFpEF patients. (55, 56). Furthermore, CRP is correlated with LV end-diastolic pressure, and CRP is associated with asymptomatic diastolic dysfunction in MetS (57). In patients with HFpEF, LV biopsies showed evidence of increased VCAM expression, increased CD3^+^, CD11^+^ and CD45^+^ leukocytes, and increased TGFβ1 expression, suggesting a link between myocardial inflammation and HFpEF (58). In the heart, inflammation can induce endothelial cell death and dysfunction leading to microvascular rarefaction and disruption of endothelial-myocyte communication with decreased NO resulting in stiff, hypertrophied cardiac myocytes (59), as well as recruitment of inflammatory monocytes and macrophages that can induce interstitial fibrosis (58, 59). Here, we found an increased number of CD64^+^ macrophages in male WT hearts on the HFpEF-MetS diet, with a 2-fold further increase in CD64^+^ macrophages in male Ffar4KO hearts, supporting the assertion that inflammation drives HFpEF. However, it remains to be determined if the increased number of CD64^+^ macrophages observed in WT hearts and the even greater increase observed in Ffar4KO hearts was causative to the observed worsened ventricular remodeling in response to MetS.

Here, we also report the first data demonstrating that loss of Ffar4 reduced cardiac 18-HEPE levels *in vivo*, supporting our prior *in vitro* data from cardiac myocytes indicating that activation of Ffar4 increased 18-HEPE levels (33). Furthermore, loss of Ffar4 increased 12-HETE levels, increasing the 12-HETE/18-HEPE ratio, and suggesting a more pro-inflammatory state in the male Ffar4KO heart both at baseline and in response to the HFpEF-MetS diet. In the heart, prior studies indicate that 12-HETE worsens ischemic injury, induces maladaptive hypertrophy, and worsens HF (60), and a recent study indicated that 12-HETE specifically worsens HFpEF remodeling (44). Ultimately, the increased 12-HETE/18-HEPE ratio in male Ffar4KO hearts was associated with an altered immune response, with increased CD64^+^ macrophage numbers, which correlated with worsened HFpEF remodeling. This suggests a novel paradigm in which one GPCR, Ffar4, functions in a feed-forward mechanism to directly (18-HEPE, ChemR23) or indirectly (12-HETE, GPR31) regulate downstream GPCR agonists to attenuate the cardiac inflammatory response to cardiometabolic disease.

Recently, we demonstrated that in adult cardiac myocytes, Ffar4-cPLA2α signaling directly and uniquely increased synthesis of 18-HEPE (33), which helps to explain the reduced levels of 18-HEPE in HDL and hearts of Ffar4KO mice both at baseline and in response to the HFpEF-MetS diet. Increased levels of 18-HEPE, secondary to EPA-supplementation or in fat-1-transgenic mice that convert ω6- to ω3-PUFAs, are associated with prevention of atherosclerosis and pathologic ventricular remodeling post-TAC (61, 62). Furthermore, 18-HEPE directly inhibits cardiac myocyte death (33), and exogenous, unesterified 18-HEPE prevents post-TAC remodeling (62). Collectively, these findings suggest that 18-HEPE is cardioprotective, but its mechanism of action remains unclear. To date, no receptor has been identified for 18-HEPE, however, 18-HEPE is the precursor for E-series resolvins (RvE), which signal through ChemR23. ChemR23 is expressed in macrophages (63), smooth muscle cells (61), endothelial cells (64), and adipocytes (65), and based on direct effects of 18-HEPE to prevent CM cell death (33) and RvE1 infusion to attenuate post-infarction remodeling (66), we speculate that ChemR23 might also be expressed in cardiac myocytes. Interestingly, ChemR23 binds to two entirely different ligands; the peptide chemerin, a macrophage chemoattractant (67), and E-series resolvins that are derived from EPA/18-HEPE (45). In macrophages, ChemR23 expression is restricted to naïve and M1-like macrophages, which respond to chemerin produced in inflamed tissue to recruit macrophages, while RvE1 promotes repolarization of M1-like macrophages towards a more resolving M2-like phenotype (68). In a mouse atherosclerosis model, EPA supplementation increased 18-HEPE levels and prevented atherosclerosis progression. However, systemic deletion of ChemR23 in this context also increased macrophage uptake of oxidized LDL, reduced phagocytosis, and increased plaque size (61). However, a separate study seemed to reach the opposite conclusion (69), which might reflect differences in the balance of the two ChemR23 ligands. In total, the reduced levels of 18-HEPE in male Ffar4KO mice suggests the potential for more pro-inflammatory signaling through macrophage ChemR23.

Positive results with icosapent ethyl (EPA) in the Reduction of Cardiovascular Events with EPA-Intervention Trial (REDUCE-IT) (70) and the Effect of Vascepa in Improving Coronary Atherosclerosis in People with High Triglycerides Taking Statin Therapy Trial (EVAPORATE) (71) to improve cardiovascular outcomes have renewed interest in the mechanistic basis for EPA-mediated cardioprotection. Following the original identification of Ffar4 (GPR120) as a receptor for long-chain fatty acids (14), there has been considerable interest in the idea that Ffar4 mediates these protective effects. However, detailed *in vitro* studies of Ffar4 pharmacology suggest that in general, PUFAs, including ω3-PUFAs (EPA) and ω6-PUFAs (AA), have relatively similar efficacy and potency, and are not biased agonists (12). Assuming this is correct, it presents a conundrum in explaining the beneficial effects of ω3-PUFAs versus ω6-PUFAs in terms of Ffar4 ligand binding or activation of immediate downstream signaling pathways. Adding to this complexity is the idea that ω3-PUFAs and EPA in particular have several proposed mechanisms of action including receptor mediated signaling through Ffar1/4 and Peroxisome Proliferator-Activated Receptors (PPARs), but also production of oxylipins (e.g. 18-HEPE), and direct effects on membrane structure (72). Our current results and previous work (33) indicate that Ffar4-cPLA2α signaling in cardiac myocytes shows surprising specificity to induce the production of 18-HEPE. Given that cPLA2α does not have a reading function and will cleave whichever PUFA is found in the sn-2 position of membrane phospholipids, it is logical to suggest that altering ω3/ω6 levels in membrane phospholipids through dietary supplementation could alter production of their cognate oxylipins. This concept is also consistent with our assertion that the mechanism by which Ffar4 prevents HFpEF remodeling is by controlling pro/anti-inflammatory oxylipin balance to attenuate inflammation.

In female Ffar4KO mice, the HFpEF-Mets diet induced a greater accumulation of fat, but surprisingly, this did not translate to worsened cardiac outcomes. If anything, female Ffar4KO hearts showed a trend towards less diastolic dysfunction. Although there is some controversy, more studies seem to suggest that loss of Ffar4 worsens metabolic outcomes with little effect on weight gain in mice challenged with HFD (19–22). Interestingly, none of these prior studies examined females. Furthermore, we specifically assessed cardiometabolic disease, by including a hypertensive challenge with the HFD, while none of these prior studies measured cardiac outcomes. Nonetheless, our results indicate that loss of Ffar4 induced 15% more weight gain over 20 weeks in female mice, suggesting that Ffar4 attenuates obesity in females. This finding might inform previous studies in which the examination of the Ffar4 R270H polymorphism and obesity in humans has produced conflicting results (20, 25). Additionally, in the context of human disease, HFpEF patients tend to be older and female (2), a population in which all HF is more prevalent (53). In our HFpEF-MetS model, female WT mice developed slightly less diastolic dysfunction, similar to previous reports suggesting female mice are protected from HFpEF (73), a seeming discrepancy with humans. Further, despite the increased weight gain, female Ffar4KO mice showed no worsening of diastolic dysfunction relative to female WT mice. One possible explanation for the sex-based difference between humans and mice is that this might simply reflect a higher survival rate of human females with cardiovascular disease who become more susceptible to HFpEF with age.

In conclusion, we demonstrate for the first time that Ffar4 attenuates cardiometabolic disease, and present novel mechanistic insight into how Ffar4 modulates oxylipin levels to attenuate inflammation and prevent HFpEF remodeling. Furthermore, we suggest a plausible mechanism through Ffar4-cPLA2α mediated EPA-derived oxylipin synthesis that sheds new light on the basis of EPA-mediated cardioprotection. Finally, based on the experimental success of synthetic Ffar4 agonist to attenuate metabolic disease (26–28), we suggest that Ffar4 might be a novel therapeutic target in cardiometabolic disease.

## Acknowledgements

The authors acknowledge the University Imaging Centers for support on echocardiography.

## Funding Sources

This work was supported by NIH HLR01130099 (TDO and GCS) and NIH HLR01152215 (TDO and GCS), Minnesota Obesity Prevention Training Program T32 NIH Grant 1T32DK083250– 01A1 (KAM), NIH Post-doctoral Fellowship F32HL152523 (MZ), American Heart Association (AHA) Grant CDA855022 (JWW) and AHA predoctoral fellowship #903380 (MTP)

## Disclosures

None

## Author Contributions

Conceptualization: N.Z., K.A.M., B.A.H., M.Z., J.W.W., G.C.S., T.D.O.

Methodology: N.Z., K.A.M., B.A.H., M.Z., B.M.W., M.T.P., C.L.H., J.W.W., G.C.S., T.D.O.

Formal analysis: N.Z., K.A.M., B.A.H., M.Z., B.M.W., M.T.P., C.L.H., J.W.W., G.C.S., T.D.O.

Investigation: N.Z., K.A.M., B.A.H., M.Z., D.J.G, B.M.W., J.M., M.T.P., D.A.O., C.L.H., J.W.W., G.C.S., T.D.O.

Writing-Original Draft: N.Z., K.A.M. B.A.H., G.C.S., and T.D.O.

Writing-Review & Editing: N.Z., K.A.M., B.A.H., C.L.H., J.W.W., G.C.S., and T.D.O.

Visualization: K.A.M., B.A.H., B.C.J, Q.S.W., G.C.S., T.D.O.

Supervision: G.C.S. and T.D.O.

Funding Acquisition: G.C.S. and T.D.O.

## Notes

### Competing Interest Statement

The authors have declared no competing interest.

