## Supplemental data for "Free fatty acid receptor 4 (FFAR4) regulates cardiac oxylipin balance to promote inflammation resolution in a model of heart failure preserved ejection fraction secondary to metabolic syndrome"

**Supplemental Table 1A:** DYETS #104607 Custom Control Diet Composition

| Ingredient | kcal/g | g/kg | kcal/kg | %kcal |
|---|---|---|---|---|
| Casein | 3.58 | 200 | 716 | 19.74 |
| L-Cystine | 4 | 3 | 12 | 0.33 |
| Sucrose | 4 | 100 | 400 | 11.03 |
| Cornstarch | 3.6 | 414.5 | 1492.2 | 41.14 |
| Dyetrose | 3.8 | 140 | 532 | 14.67 |
| Corn oil | 9 | 25 | 225 | 6.20 |
| Lard | 9 | 20 | 180 | 4.96 |
| Cellulose | 0 | 50 | 0 | 0.00 |
| Mineral Mix #210025 | 0.88 | 35 | 30.8 | 0.85 |
| Vitamin Mix #310025 | 3.87 | 10 | 38.7 | 1.07 |
| Choline Bitartrate | 0 | 2.5 | 0 | 0.00 |
| **Total** | | **1000** | **3626.7** | **100** |

**Supplemental Table 1B:** DYETS #104608 Custom High Fat Diet Composition

| Ingredient | kcal/g | g/kg | kcal/kg | %kcal |
|---|---|---|---|---|
| Casein | 3.58 | 200 | 716 | 15.56 |
| L-Cystine | 4 | 3 | 12 | 0.26 |
| Sucrose | 4 | 350 | 1400 | ***30.42*** |
| Cornstarch | 3.6 | 134.5 | 484.2 | 10.52 |
| Corn oil | 9 | 25 | 225 | 4.89 |
| Lard | 9 | 190 | 1710 | ***37.15*** |
| Cellulose | 0 | 50 | 0 | 0.00 |
| Mineral Mix #200000 | 0.47 | 35 | 16.45 | 0.36 |
| Vitamin Mix #310035 | 3.92 | 10 | 39.2 | 0.85 |
| Calcium Carbonate | 0 | 2.5 | 0 | 0.00 |
| **Total** | | **1000** | **4602.85** | **100** |

**Supplemental Table 2A:** Metabolic parameters in **male** mice after 20 weeks on diet

| **Metabolic Parameter** | **WT Cont (15)** | **KO Cont (20)** | **WT HFpEF (17)** | **KO HFpEF (20)** | **ANOVA** Primary | Diet | Genotype |
|---|---|---|---|---|---|---|---|
| BW (g) | 32.2 (31.1, 33.3) | 31.4 (30.4, 32.5) | 46.9 (44.4, 49.3) | 48.8 (47.4, 50.2) | 0.0785 | <0.0001 | 0.4293 |
| Fat mass (g) | 4.5 (3.1, 5.8) | 3.9 (2.6, 5.2) | 17.7 (16.0, 19.0) | 19.1 (18.0, 20.0) | 0.1199 | <0.0001 | 0.5028 |
| Lean mass (g) | 27.0 (25.7, 28.4) | 26.8 (26.1, 27.5) | 28.8 (27.4, 30.1) | 29.4 (28.6, 30.2) | 0.3815 | <0.0001 | 0.7032 |
| Adiposity index | 0.2 (0.1, 0.2) | 0.1 (0.1, 0.2) | 0.6 (0.6, 0.7) | 0.7 (0.6,0.7) | 0.1600 | <0.0001 | 0.7116 |
| Plasma TG (nM) | 35.0 (29.6, 40.3) | 48.8 (43.7, 54.0) | 79.2 (74.1, 84.3) | 93.1 (87.9, 98.3) | 0.0160 | <0.0001 | 0.0044 |
| Plasma HDL (nM) | 41.1 (37.8, 44.4) | 44.2 (403., 48.2) | 60.6 (57.5, 64.0) | 72.3 (68.1, 76.3) | 0.0124 | <0.0001 | 0.0004 |
| SBP (mmHg) | 103.6 (100.2, 107.1) | 102.4 (99.1, 105.7) | 131.8 (127.3, 136.2) | 134.0 (129.4, 138.6) | 0.3741 | <0.0001 | 0.8063 |
| DBP (mmHg) | 77.1 (73.5, 80.8) | 75.3 (71.9, 78.7) | 96.5 (92.7, 100.3) | 103.1 (98.7, 107.4) | 0.0266 | <0.0001 | 0.2099 |
| MAP (mmHg) | 85.6 (82.1, 89.2) | 84.0 80.7, 87.3) | 107.9 (104.1, 111.8) | 113.0 (108.7, 117.4) | 0.0716 | <0.0001 | 0.3485 |

**Supplemental Table 2B:** Metabolic parameters in **female** mice after 20 weeks on diet

| **Metabolic Parameter** | **WT Cont (17)** | **KO Cont (18)** | **WT HFpEF (16)** | **KO HFpEF (17)** | **ANOVA** Primary | Diet | Genotype |
|---|---|---|---|---|---|---|---|
| BW (g) | 21.9 (21.3, 22.4) | 22.9 (22.0, 23.7) | 30.9 (29.0, 32.7) | 35.5 (32.7, 38.2) | 0.0306 | <0.0001 | 0.0010 |
| Fat mass (g) | 2.1 (1.8, 2.4) | 2.4 (2.0, 2.8) | 10.5 (8.9, 12.0) | 14.6 (12.4, 16.8) | 0.0043 | <0.0001 | 0.0008 |
| Lean mass (g) | 18.8 (18.4, 19.2) | 19.5 (18.9, 20.1) | 19.8 (19.3, 20.3) | 20.5 (19.5, 21.5) | 0.9645 | 0.0020 | 0.0323 |
| Adiposity index | 0.1 (0.09,0.1) | 0.1 (0.1,0.1) | 0.5 (0.5, 0.6) | 0.7 (0.6, 0.8) | 0.0039 | <0.0001 | 0.0008 |
| Plasma TG (nM) | 39.7 (34.8, 45.4) | 53.6 (48.8, 58.3) | 72.5 (67.4, 77.6) | 86.4 (81.4, 91.4) | 0.4144 | <0.0001 | <0.0001 |
| Plasma HDL (nM) | 35.8 (32.6, 39.0) | 39.0 (35.1, 42.9) | 55.3 (52.1, 58.6) | 67.0 (62.9, 71.1) | 0.2429 | <0.0001 | 0.0167 |
| SBP (mmHg) | 107.0 (102.7, 111.6) | 103.7 (100.1, 107.3) | 139.7 (134.5, 144.9) | 132.0 (128.0, 136.0) | 0.2908 | <0.0001 | 0.0076 |
| DBP (mmHg) | 80.2 (76.2, 84.3) | 77.2 (73.4, 81.1) | 108.7 (103.6, 113.8) | 99.9 (94.9, 105.0) | 0.1836 | <0.0001 | 0.0077 |
| MAP (mmHg) | 88.9 (84.7, 93.0) | 85.7 (82.1, 89.4) | 114.6 (106.6, 122.5) | 110.3 (105.7, 115.0) | 0.8283 | <0.0001 | 0.1408 |

Data are presented as Mean ± standard error for the number of mice indicated in parentheses. TG, triglyceride; HDL, high density lipoprotein; SBP, systolic blood pressure; DBP, diastolic blood pressure; MAP; mean arterial pressure. ANOVA denotes the results of the two-way ANOVA with the *P* values shown for the primary interaction as well as the effects of diet or genotype.

**Supplemental Table 3A:** Cardiac function in **male** mice after 20 weeks on diet

| Cardiac function | WT Cont (15) | KO Cont (20) | WT HFpEF (17) | KO HFpEF (20) | ANOVA primary | Diet | Genotype |
|---|---|---|---|---|---|---|---|
| EF (%) | 50.5 (45.4, 55.7) | 53.8 (48.8, 58.9) | 55.7 (50.8, 60.5) | 47.6 (42.5, 52.8) | 0.0236 | 0.8269 | 0.3424 |
| SV (µl) | 38.3 (35.3, 41.4) | 39.9 (34.6, 45.2) | 39.5 (35.1, 44.0) | 34.5 (30.8, 38.3) | 0.1233 | 0.3181 | 0.4211 |
| FS (%) | 25.7 (22.5, 28.9) | 27.9 (24.7, 31.1) | 28.9 (25.8, 32.1) | 23.9 ( 20.9, 27.0) | 0.0201 | 0.8250 | 0.3525 |
| EDV (µl) | 77.3 (70.3, 84.2) | 74.6 (66.6, 82.5) | 72.1 (63.8, 80.4) | 73.9 (65.9, 81.8) | 0.5548 | 0.4386 | 0.9047 |
| ESV (µl) | 38.9 (31.7, 46.2) | 34.7 (28.9, 40.6) | 32.6 (26.0, 39.2) | 39.3 (32.3, 46.4) | 0.0974 | 0.7920 | 0.7028 |
| CO (ml/min) | 17.4 (15.3, 19.4) | 18.6 (16.0, 21.2) | 16.5 (14.5, 18.4) | 15.2 (13.4, 16.9) | 0.2158 | 0.0391 | 0.9802 |
| IVS; s (mm) | 1.4 (1.3, 1.5) | 1.4 (1.3, 1.4) | 1.5 (1.5, 1.6) | 1.5 (1.4, 1.6) | 0.6494 | 0.0002 | 0.8005 |
| IVS; d (mm) | 1.0 (0.9, 1.1) | 1.0 (0.9, 1.0) | 1.1 (1.1, 1.2) | 1.1 (1.1, 1.2) | 0.4849 | <0.0001 | 0.8141 |
| LVID; s (mm) | 3.1 (2.9, 3.3) | 3.0 (2.8, 3.1) | 2.9 (2.6, 3.1) | 3.1 (2.9, 3.3) | 0.0592 | 0.6635 | 0.7197 |
| LVID; d (mm) | 4.2 (4.0, 4.3) | 4.1 (3.9, 4.3) | 4.0 (3.8, 4.2) | 4.1 (3.9, 4.2) | 0.5636 | 0.3368 | 0.7865 |
| LVPW; s (mm) | 1.1 (1.0, 1.2) | 1.0 (0.9, 1.1) | 1.2 (1.1, 1.3) | 1.1 (1.0, 1.2) | 0.8638 | 0.0098 | 0.1123 |
| LVPW; d (mm) | 0.8 (0.7, 0.8) | 0.7 (0.6, 0.8) | 0.9 (0.8, 0.9) | 0.9 (0.8, 0.9) | 0.3836 | <0.0001 | 0.1023 |
| LV mass corrected (mg) | 114.7 (106.8, 122.6) | 105.2 (96.0, 114.4) | 128.7 (120.1, 137.3) | 132.4 (119.9, 144.9) | 0.1763 | <0.0001 | 0.5536 |
| LVRI | 34.4 (32.5, 36.3) | 32.0 (29.9, 34.1) | 39.8 (38.4, 41.3) | 40.8 (37.4, 44.2) | 0.1568 | <0.0001 | 0.5511 |
| E/A | 1.4 (1.3, 1.5) | 1.4 (1.3, 1.5) | 1.8 (1.6, 2.0) | 2.1 (1.9, 2.4) (18) | 0.0779 | <0.0001 | 0.0415 |
| E (mm/s) | 784.2 (740.1, 828.2) | 724.6 (695., 753.5) | 748.6 (692.1, 805.2) | 795.6 (755.9, 835.3) | 0.0102 | 0.3818 | 0.7554 |
| A (mm/s) | 562.3 (528.3, 596.3) | 513.5 (482.9, 544.1) | 422.9 (377.0, 468.8) | 388.8 (344.1, 433.6) (18) | 0.6941 | <0.0001 | 0.0301 |
| E/e' | 24.9 (22.2, 27.6) | 24.7 (23.0, 26.5) (19) | 31.9 (29.3, 34.4) | 42.1 (36.7, 47.6) | 0.0036 | <0.0001 | 0.0048 |
| GLS (%) | -18.7 (-20.9, -16.5) | -22.0 (-24.41, -19.47) | -17.4 (-19.82, -14.93) | -14.2 (-16.0, -12.4) | 0.0045 | <0.0001 | 0.9918 |
| RLPSR (1/s) | 7.7 (6.1, 9.4) | 8.8 (7.3, 10.5) | 7.5 (5.7, 9.4) | 5.6 (4.7, 6.4) | 0.0463 | 0.0224 | 0.5979 |
| PAT norm. to HR (ms) | 16.1 (13.9, 18.2) (8) | 18.6 (16.9, 20.3) (6) | 14.4 (12.8, 16.0) (8) | 14.3 (13.3, 15.4) (12) | 0.0717 | 0.0002 | 0.0904 |
| HR (bpm) | 450.7 (427.1, 474.3) | 466.7 (447.5, 485.8) | 417.4 (403.1, 431.8) | 439.5 (421.7, 457.3) | 0.7359 | 0.0013 | 0.0377 |

**Supplemental Table 3B:** Cardiac function in **female** mice after 20 weeks on diet

| Cardiac function | WT Cont (17) | KO Cont (18) | WT HFpEF (16) | KO HFpEF (17) | ANOVA primary | Diet | Genotype |
|---|---|---|---|---|---|---|---|
| EF (%) | 59.31 (55.2, 63.4) | 54.6 (47.4, 61.8) | 48.4 (43.6, 53.3) (15) | 52.3 (46.4, 58.2) | 0.1183 | 0.0183 | 0.8833 |
| SV (ul) | 32.33 (30.0, 34.6) | 31.0 (27.3, 34.6) | 27.8 (25.0, 30.5) (15) | 29.3 (25.8, 32.9) | 0.3335 | 0.0438 | 0.9591 |
| FS (%) | 31.04 (28.3, 33.8) | 28.6 (23.8, 33.3) | 24.1 (21.1, 27.2) (15) | 26.8 (22.8, 30.7) | 0.1553 | 0.0179 | 0.9634 |
| EDV (ul) | 54.89 (51.5, 58.2) | 58.0 (53.5, 62.4) | 57.9 (53.6, 62.2) (15) | 56.4 (51.5, 61.2) | 0.2631 | 0.7355 | 0.7056 |
| ESV (ul) | 22.57 (19.5, 25.6) | 27.0 (21.2, 32.9) | 30.1 (25.9, 34.3) (15) | 27.0 (22.9, 31.2) | 0.0831 | 0.0810 | 0.7461 |
| CO (ml/min) | 14.4 (13.0, 15.9) | 14.8 (13.0, 16.5) | 11.9 (10.5, 13.3) (15) | 13.5 (11.9, 15.1) | 0.4240 | 0.0117 | 0.2080 |
| IVS; s (mm) | 1.3 (1.2, 1.4) | 1.3 (1.2, 1.4) | 1.3 (1.2, 1.4) (15) | 1.3 (1.2, 1.5) | 0.7024 | 0.8485 | 0.9660 |
| IVS; d (mm) | 0.9 (0.9, 1.0) | 0.9 (0.9, 1.0) | 1.0 (0.9, 1.1) (15) | 1.0 (0.9, 1.1) | 0.6444 | 0.1075 | 0.9231 |
| LVID; s (mm) | 2.5 (2.4, 2.6) | 2.6 (2.4, 2.9) | 2.2 (2.6, 3.0) (15) | 2.6 (2.5, 2.9) | 0.0983 | 0.0645 | 0.9132 |
| LVID; d (mm) | 3.6 (3.5, 3.7) | 3.7 (3.6, 3.8) | 3.7 (3.6, 3.8) (15) | 3.6 (3.5, 3.8) | 0.3370 | 0.8626 | 0.8092 |
| LVPW; s (mm) | 1.1 (1.0, 1.2) | 1.0 (0.8, 1.0) | 1.0 (0.9, 1.1) (15) | 0.9 (0.8, 1.0) | 0.3758 | 0.4224 | 0.0275 |
| LVPW; d (mm) | 0.8 (0.7, 0.8) | 0.6 (0.6, 0.7) | 0.8 (0.7, 0.8) (15) | 0.7 (0.6, 0.8) | 0.4449 | 0.2925 | 0.0367 |
| LV mass corrected (mg) | 84.4 (78.6, 90.2) | 80.6 (73.7, 87.5) | 94.7 (83.6, 105.8) (15) | 87.5 (76.7, 98.3) | 0.6819 | 0.0415 | 0.1881 |
| LVRI | 29.2 (27.3, 30.9) | 27.5 (24.9, 30.0) | 32.0 (28.9, 35.1) (15) | 30.2 (26.3, 34.0) | 0.9576 | 0.0484 | 0.2083 |
| E/A | 1.4 (1.3, 1.5) | 1.5 (1.3, 1.7) (16) | 2.5 (2.0, 3.0) (15) | 2.1 (1.7, 2.4) | 0.0505 | <0.0001 | 0.2498 |
| E (mm/s) | 680.3 (619.0, 741.6) | 672.5 (618.3, 726.7) (17) | 754.0 (710.4, 797.7) | 773.6 (717.9, 829.3) | 0.5955 | 0.0011 | 0.8201 |
| A (mm/s) | 488.4 (449.9, 527.0) | 448.8 (407.0, 490.5) | 336.0 (273.4, 398.6) (15) | 397.1 (352.8, 441.3) | 0.0256 | <0.0001 | 0.6285 |
| E/e' | 22.4 (21.0, 23.8) | 22.6 (20.6, 24.7) (17) | 37.8 (32.0, 43.7) (16) | 31.3 (27.1, 35.5) | 0.0806 | <0.0001 | 0.1057 |
| HR (bpm) | 445.0 (422.5, 467.4) | 479.5 (453.7, 505.3) | 428.3 (411.0, 445.6) (15) | 459.0 (442.1, 478.9) | 0.9082 | 0.0872 | 0.0018 |

Data are presented as Mean ± standard error for the number of mice indicated in parentheses. HR, heart rate; SV, stroke volume; EF, ejection fraction; FS, fractional shortening; EDV, end diastolic volume; ESV, end systolic volume; CO, cardiac output; IVS;s, interventricular septal thickness at systole; IVS;d, interventricular septal thickness at diastole; LVID;s, left ventricular internal diameter systole; LVID;d, left ventricular internal diameter diastole; LVPW;s, left ventricular posterior wall systole; LVPW;d, left ventricular posterior wall diastole; LVRI, left ventricular remodeling index; E/A, early mitral valve filling velocity/ late mitral valve filling velocity; E, early mitral valve filling velocity ; E/E', early mitral valve filling velocity/early mitral annular tissue velocity; GLS, global longitudinal strain; RLPSR, reverse longitudinal peak strain rare; PAT, pulmonary artery acceleration time normalized to heart rate. ANOVA denotes the results of the two-way ANOVA with the *P* values shown for the primary interaction as well as the effects of diet or genotype.

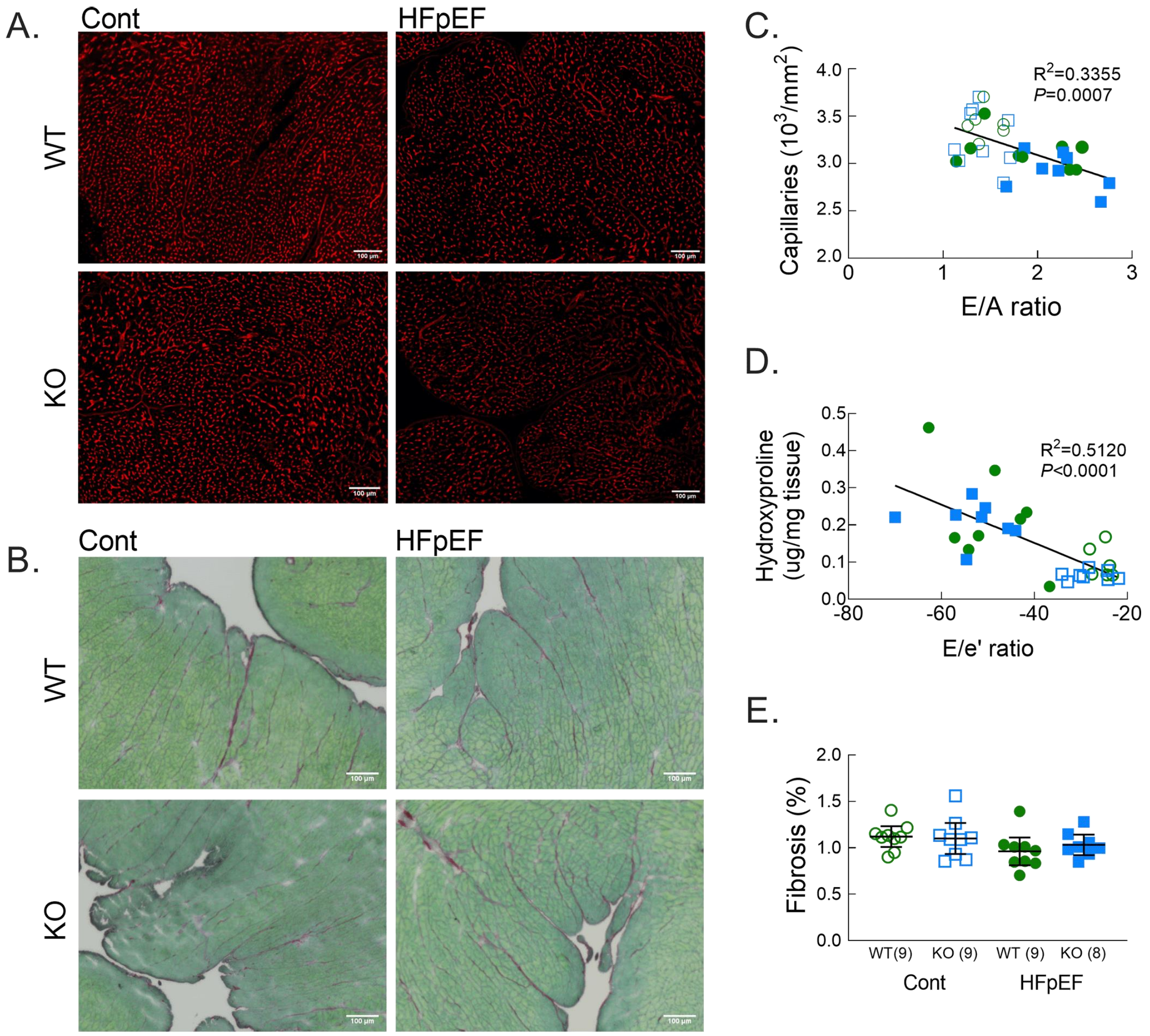


**Supplemental Figure 1**

**A.** Representative images of capillaries in LV sections from male WT and Ffar4KO mice stained with isolectin B4 (IB4) (Red). **B.** Representative images of ventricular fibrosis quantified from ventricular cross sections from male WT and Ffar4KO mice stained with Sirius red/Fast Green (SR/FG). **C.** Relationship between myocardial capillary density and diastolic function (E/A ratio) n=32, $R^2$=0.3355, P=0.0007. **D.** Relationship between ventricular fibrosis quantified by hydroxyproline content and diastolic function (E/e' ratio) n=30, $R^2$=0.5120, P<0.0001. **E.** Ventricular fibrosis quantified by SR/FG staining.
